# Projected climate change will reduce habitat suitability for bumble bees in the Pacific Northwest

**DOI:** 10.1101/610071

**Authors:** Jonathan B. Koch, Chris Looney, Brandon Hopkins, Elinor M. Lichtenberg, Walter S. Sheppard, James P. Strange

**Affiliations:** Department of Biology & Ecology Center, Utah State University, 5305 Old Main Hill, Logan, Utah, 84322; United States Department of Agriculture-Agricultural Research Services, Pollinating Insects-Biology, Management, and Systematics Research Laboratory, 5305 Old Main Hill, Logan, Utah, 84322; Plant Protection Division, Washington State Department of Agriculture, Olympia, WA 98504; Department of Entomology, Washington State University, Pullman, WA 99164; Department of Integrative Biology, The University of Texas at Austin, Austin, TX 78712

## Abstract

Global climate change is the greatest environmental challenge of the modern era. The impacts of climate change are increasingly well understood, and have already begun to materialize across diverse ecosystems and organisms. Bumble bees (*Bombus*) are suspected to be highly sensitive to climate change as they are predominately adapted to temperate and alpine environments. In this study, we determine which bumble bee species are most vulnerable to climate change in the Pacific Northwest. The Pacific Northwest is a topographically complex landscape that is punctuated by two major mountain ranges and a labyrinth of offshore islands in the Salish Sea. Using standardized survey methods, our study documents the occurrence of 15 bumble bee species across 23 field sites in seven federal parks, historical sites, and monuments. Our results show that bumble bee community richness and diversity increases along an altitude gradient in these protected areas. Furthermore, NMDS analysis reveals that high altitude environments are composed of a unique group of bumble bee species relative to low altitude environments. Finally, based on an analysis of species distributions models that aggregate bioclimatic data from global circulation climate models with preserved specimen records, we discover that 80% of the bumble bee species detected in our survey are poised to undergo habitat suitability (HS) loss within the next 50 years. Species primarily found in high altitude environments namely *B. vandykei, B. sylvicola*, and *B. bifarius* are projected to incur a mean HS loss of 63%, 59%, and 30% within the federally protected areas, respectively. While the implementation of climate change policies continue to be a significant challenge, the development of mitigation strategies to conserve the most vulnerable species may be a tractable option for land managers and stakeholders of protected areas. Our study meets this need by identifying which species and communities are most sensitive to climate change.

## Introduction

Pollinator communities worldwide are undergoing dramatic changes in both abundance and composition that may put pollination service at risk in many terrestrial ecosystems [1]. These changes may not solely be unidirectional declines in species abundance, but can manifest as shifts in geographic range, increases in abundance where new habitat is formed, or shifts in phenology [2–4]. To date, documented changes in pollinator communities have been attributed to many factors, including pathogen outbreaks, pesticides, climate change, introduced species, and land-use change [4–9]. Identifying the factors affecting pollinator communities can be challenging as most strategies investigate distinct taxonomic groups (guilds) [10], or attempt to isolate specific threats and measure a single species’ responses to the threat in question [2–4,11]. However, given the negative impacts of rapid global change [1,5], it is imperative to identify which pollinator species out of a guild might be most vulnerable to a specific environmental impact. Identifying the most at-risk or vulnerable species within a guild might allow for a more effective approach to management and threat mitigation [12,13].

In montane regions, some pollinators are predicted to follow plant distributional shifts up slope as climates warm [14,15], but where species are already restricted to high altitude habitat it is unclear if they can adapt *in situ* [16–18]. Climate change is an emerging threat to pollinators, yet it remains poorly studied because isolating the effects of climate from other potential factors is difficult [4,19]. There is a global consensus among scientists that the economic activities associated with human population growth have significantly influenced climate patterns by increasing greenhouse gas emissions, namely carbon dioxide (CO_2_), methane (CH_4_), and nitrous oxide (N_2_O) since the industrial revolution [20]. Bee phenology derived from museum records already demonstrates earlier springtime activity of bees in the northeastern US, correlated with climate warming over the past century [4]. Furthermore, the reduction of suitable habitat due to climate change is suspected to shift bumble bee distributions upslope [15], a phenomenon observed in diversity of organisms [17,18]. Miller-Struttmann et al. [16] found that alpine bumble bee species have experienced rapid evolutionary change in the length of their proboscis due to the decline of floral resources in montane regions in Colorado. Studies like Miller-Struttmann et al. thereby suggest that species that are adapted to alpine environments might be exposed to greater evolutionary pressures in the next 50 years due to climate change [17,21–24]. Under currently projected climate models, bumble bee distributions are predicted to shift to higher latitudes in the cases where habitat suitability gains in altitude are limited [2,3,8].

Bumble bees (*Bombus*) are a predominantly temperate-adapted genus of primitively eusocial bee (Hymenoptera: Apidae) that are dependent on a variety of floral resources for pollen and nectar [5]. There are more than 250 different bumble bee species worldwide, 30 of which are distributed in the western US [25,26]. They are important pollinators of wild flowering plants especially in montane and alpine environments. The US Pacific Northwest is rich in wildflower and bumble bee diversity, largely in part to the environmental heterogeneity resulting from the region’s complex topography [26,27]. The topography of the Pacific Northwest is hypothesized to have significantly influenced patterns of population genetic diversity [27,28], with some protected mountain and island regions lending themselves to uncommon phenotypes of certain bumble bee species [29,30]. In the Pacific Northwest *B. occidentalis* is known to be at risk for decline due to pathogens, while *B. vosnesenskii*, may be expanding in range [10,31]. Several other species are also suspected of undergoing changes in range or abundance in the region, yet empirical data is currently lacking [10,32,33]. While a high richness of bumble bee species is found in the Pacific Northwest [26], they are threatened by the effects of projected climate change in the region. It is estimated that over the next century, the region will incur rates of warming by up to 1°C per decade and a 1–2% increase in annual precipitation, likely facilitating wetter autumns and drier summers and winters [34].

There is a critical need to estimate the effects of projected climate change on bumble bee communities in the Pacific Northwest. While domestic and international policy will be the key factor in mitigating the effects of climate change, managers of protected areas in the US may begin to develop management and prioritization strategies for species that are most vulnerable to the effects of climate change [12,35]. The US National Parks found in the Pacific Northwest are situated across an altitude gradient that allows for an investigation on community composition and turnover of bumble bee pollinators and an assessment of the impacts of projected climate change on bumble bee habitat suitability. In this study, we aim to answer the following questions: 1) What is the relationship between species diversity/richness across an altitude gradient? 2) Is bumble bee community composition predicted by their distribution across an altitude gradient? And 3) Which species will experience significant gains/losses in habitat suitability in the Pacific Northwest national parks based on projected climate change scenarios? To answer these questions, we surveyed bumble bee communities to estimate species richness and diversity. We then constructed species distribution models (SDMs) to estimate habitat suitability (HS) for the bumble bees distributed in the region by combining georeferenced museum records with bioclimatic data. Finally, we projected the SDMs to future climate scenarios to estimate HS change. Characterizing bumble bee community composition and projecting HS change in the Pacific Northwest will provide park management with information on which bumble bee communities and species are most vulnerable to climate change.

## Materials and Methods

### Field Survey

In the summers of 2013 and 2014 we visited 23 field sites in seven US National Parks in the Pacific Northwest to survey bumble bees (Fig 1; Table 1). In Olympic National Park (OLYM) and Mount Rainier National Park, two transects were surveyed across an altitude gradient, while North Cascades National Park (NOCA) had one transect surveyed. In Ebey’s Landing National Historical Reserve (EBLA), Lewis and Clark National Historical Park (LEWI), and Fort Vancouver National Historic Site (FOVA) one site at each park was surveyed. In San Juan Islands National Historical Park (SAJH), two sites were surveyed. We did not survey bumble bees across an altitude gradient in EBLA, LEWI, FOVA, and SAJH as they are near sea level. In NOCA and OLYM, we revisited some sites surveyed in a previous study [10].

**Table 1.** Bumble bee field sites and abundances across and adjacent to US National Parks in the Pacific Northwest.

| Site | Altitude<br>(m) | National<br>Park | Latitude | Longitude | Abundance |
| --- | --- | --- | --- | --- | --- |
| Ebey's Landing | 20 | Ebey's<br>Landing | 48.1933 | -122.7096 | 35 |
| Fort Vancouver | 12 | Fort<br>Vancouver | 45.6236 | -122.6615 | 55 |
| Lewis & Clark NP | 2 | Lewis &<br>Clark | 46.1175 | -123.8752 | 33 |
| Lower Palisades Lake | 1804 | Mt. Rainier | 46.9542 | -121.5924 | 5 |
| near Sunrise Meadows | 1907 | Mt. Rainier | 46.9136 | -121.6222 | 8 |
| Paradise Meadows | 1603 | Mt. Rainier | 46.7863 | -121.7399 | 48 |
| Snow Lake | 1458 | Mt. Rainier | 46.7655 | -121.7057 | 44 |
| Upper Crystal Lake | 1784 | Mt. Rainier | 46.9057 | -121.5094 | 53 |
| Upper Palisades Lake | 1804 | Mt. Rainier | 46.9493 | -121.5923 | 30 |
| West Side Road | 878 | Mt. Rainier | 46.7794 | -121.8847 | 50 |
| Sahale Arm Trail | 1875 | North<br>Cascade | 48.4713 | -121.5188 | 44 |
| Sibley Creek | 419 | North<br>Cascade | 48.5122 | -121.2507 | 36 |
| Cascade Pass | 1638 | North<br>Cascades | 48.468 | -121.0596 | 38 |
| Crescent Lake, East Beach | 209 | Olympic | 48.086 | -123.7429 | 17 |
| Heart Lake | 1460 | Olympic | 47.9109 | -123.733 | 28 |
| Lower Bridge Creek<br>Campsite | 1163 | Olympic | 47.9241 | -123.7334 | 42 |
| Lower Royal Basin | 1421 | Olympic | 47.8391 | -123.2113 | 41 |
| Royal Basin Parking Lot | 1170 | Olympic | 47.8776 | -123.0039 | 5 |
| Royal Basin Ranger Station | 1564 | Olympic | 47.8331 | -123.2112 | 25 |
| Royal Basin Trail | 1177 | Olympic | 47.8592 | -123.2028 | 2 |
| Sandpoint Loop | 12 | Olympic | 48.1544 | -124.69 | 13 |
| American Camp | 38 | San Juan<br>Island | 48.4612 | -123.0221 | 61 |
| English Camp | 1 | San Juan<br>Island | 48.5862 | -123.1502 | 60 |

**Fig 1.**
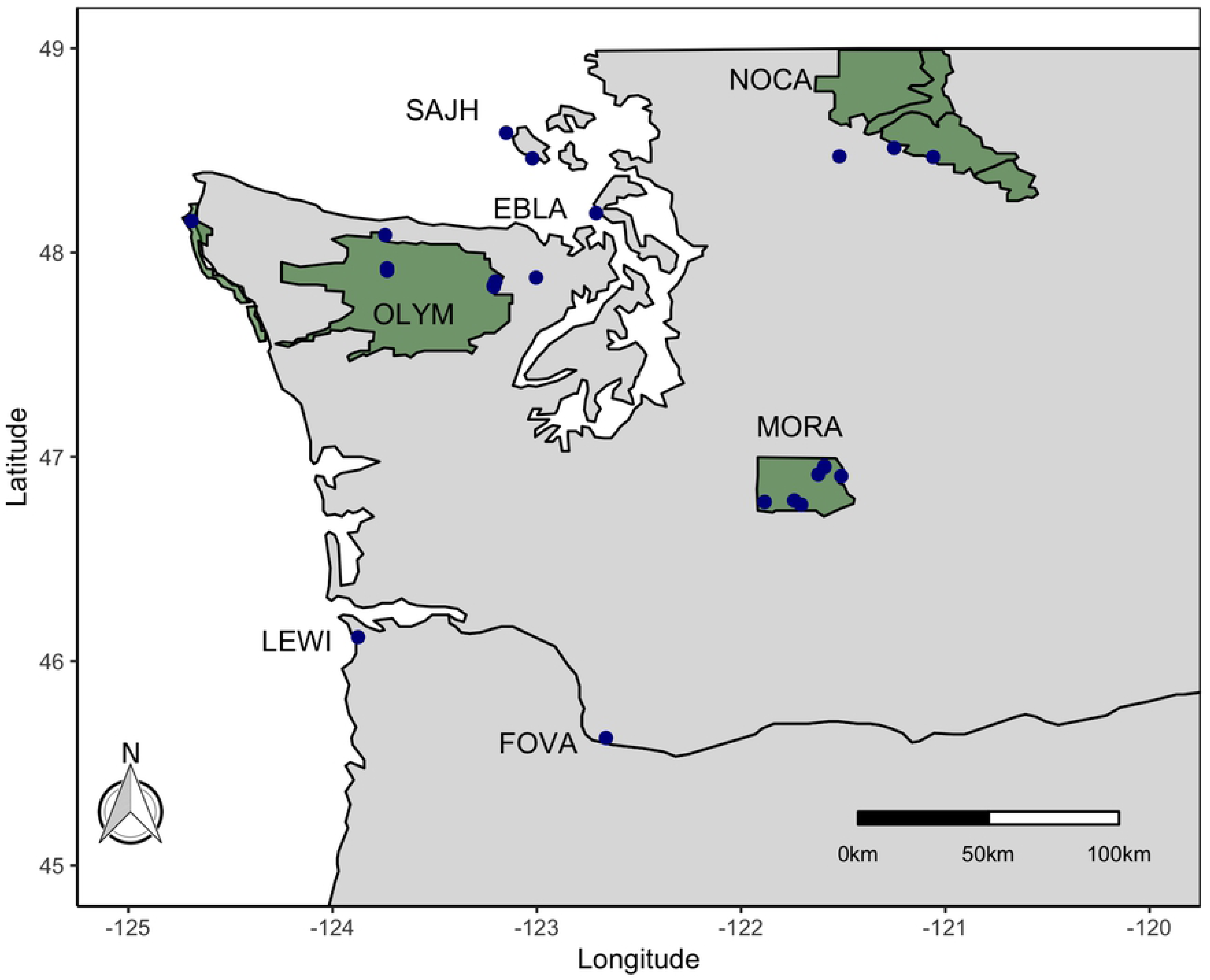
Distribution of field sites surveyed for bumble bees across and adjacent to US National Parks in the Pacific Northwest. National Parks are demarcated by large green polygons, and field site are demarcated by dark blue points. OLYM=Olympic National Park, MORA=Mount Rainier National Park, NOCA=North Cascades National Park, EBLA=Ebey’s Landing National Historical Reserve, LEWI= Lewis and Clark National Historical Park, FOVA= Fort Vancouver National Historic Site, SAJH= San Juan Islands National Historical Park.

Sites were surveyed by teams of individuals using standardized net collections of bumble bees at plots of approximately 0.5 ha. Sites varied in floral density and accessibility for off trail movement. To standardize sampling effort, surveys were timed and collections were numerically synchronized to 1.5 collector hours per site when feasible. Collectors surveyed with entomological nets (30 cm diameter) and collected bumble bees foraging on flowers directly into 20 mL plastic vials. The vials were placed on ice for 10-15 minutes until surveys were complete and the bees were immobilized by the cold. Upon completion of the survey period, the bumble bees were sexed and preliminarily identified to species using regional field guides [26,36]. While the specimens were immobilized, we non-lethally sampled DNA from the bumble bees by removing a mid-leg from each individual [27]. The mid-legs were individually stored in 95% ethanol for DNA analysis to verify the species identity. At each site, a worker and male of each captured species were sacrificed and retained as voucher specimens. All queens were released after legs were sampled.

During the survey period we recorded floral hosts to each specimen and collected pertinent environmental data from each site. Each survey event was assigned a unique locality description and georeferenced with a Garmin GPSmap 60CS. We recorded temperature (°C), relative humidity (%), and wind speed (kph) data with a Kestrel 4000 Pocket Weather Tracker. Voucher specimens were pinned and assigned a unique barcode ID, and curated into the USDA-ARS National Pollinating Insect Collection (NPIC) in Logan, UT (Table S2). Genotyped individuals were given a unique ID, an NPS accession number, and included in the NPIC database. The data is stored digitally in a relational database at the NPIC, and is also readily available on the National Park Research Permit and Reporting System website (https://irma.nps.gov/rprs/IAR/Profile/103061).

### Community analysis

We estimated species richness and diversity across the bumble bee communities using individual-based rarefaction. Species diversity was estimated with the inverse Simpson’s D index (1/D). We tested for a correlation among species richness, diversity, and altitude with a Spearman Rank-Order Correlation test. Because of unequal sample size across field sites, we used rarefaction to estimated species richness and diversity [37]. However, we first removed four field sites from the rarefaction analysis as we detected less than 10 bumble bee individuals from the survey attempt. The remaining 19 field sites were rarefied to *n* = 10 with the function *rarefy()*. Pairwise community dissimilarity was examined using the Bray-Curtis index in a non-metric dimensional scaling (NMDS) analysis. We then used the function *envfit()* to fit the altitude variable to the ordination results from the NMDS analysis, with 999 permutations. Projecting the ordination points onto the altitude variable (*i.e.,* environmental vector) allows us to test for a correlation between the two values. Rarefaction, NMDS, and the *envfit()* function are available in the vegan 2.5.3 library in R [38].

### Species distribution modelling

We queried the Global Biodiversity Information Facility website (GBIF) (http://gbif.org) for bumble bee specimen records to be used in constructing SDMs. We limited our query to only include records that were “Preserved Specimens” to maximize the probability that the specimens were identified using a taxonomic key or a voucher collection. To estimate habitat suitability (HS), SDMs were constructed under the principle of maximum entropy with MaxEnt v3.4.0 [39,40]. The algorithm in MaxEnt uses presence-only georeferenced spatial data and random background points sampled from the study extent to estimate the distribution of the species that is closest to uniform (=maximum entropy) under the suite of independent variables (*i.e.,* bioclimatic variables) supplied to the model [41]. HS is constrained between 0 and 1, where values closer to 0 represent low HS for the target bumble bee species, and values closer to 1 represent high HS for the target bumble bee species. Specifically, HS is a measure of how suitable an area unit is based on the known distribution (specimen occurrence record) of the target species and supplied bioclimatic variables.

We approximated HS for 15 bumble bee species distributed in the parks by aggregating occurrence records with a suite of 19 bioclimatic variables representing contemporary conditions (1950-2000) from the WorldClim v1.4 Bioclim database. The bioclimatic variables investigated included: BIO1 = annual mean temperature, BIO2 = mean diurnal range (mean of monthly (maximum temp - minimum temp)), BIO3 = isothermality (BIO2/BIO7) (* 100), BIO4 = temperature seasonality (standard deviation *100), BIO5 = maximum temperature of warmest month, BIO6 = minimum temperature of coldest month, BIO7 = temperature annual range (BIO5-BIO6), BIO8 = mean temperature of wettest quarter, BIO9 = mean temperature of driest quarter, BIO10 = mean temperature of warmest quarter, BIO11 = mean temperature of coldest quarter, BIO12 = annual precipitation, BIO13 = precipitation of wettest month, BIO14 = precipitation of driest month, BIO15 = precipitation seasonality (coefficient of variation), BIO16 = precipitation of wettest quarter, BIO17 = precipitation of driest quarter, BIO18 = precipitation of warmest quarter, BIO19 = precipitation of coldest quarter. Bioclimatic variables were downloaded at a spatial resolution of 2.5 arc minutes (~5 km^2^) and clipped to the spatial extent of the western US (Northernmost latitude: 49 Southernmost latitude: 30, Easternmost longitude: −100, Westernmost longitude: −125; Geographic Projection: WGS1984) (http://worldclim.org) [42].

To reduce model complexity, we examined the relationship between the 19 continuous bioclimatic variables with a pairwise Pearson correlation coefficient (*r*) test across all 15 species. From each pairwise correlation coefficient estimate, we randomly retained only one variable for the final model if *r* ≥ 0.80. If more than two specimen records fell within a raster pixel of the bioclimatic data, only one specimen record was retained for the final SDM. With MaxEnt, we constructed the SDMs using the default parameters of the program to generate a complementary log-log transformation (cloglog) to produce an estimate of habitat suitability averaged over 100 replicates with a subsampling scheme to evaluate model performance (75% train, 25% test) [40]. Models were evaluated with the area under the curve statistic (AUC). Values of AUC of 0.5 connote performance no better than random, and values < 0.5 worse than random. Thus, AUC > 0.5 is the cutoff for “good” models [39]. Each variable was evaluated for its relative importance to each species’ SDM by estimating percent contribution. In each iteration of the training algorithm, the increase in regularized gain is added to the contribution of the corresponding variable. Conversely, the regularized training gain is subtracted from the contribution of the corresponding variable if the change to the absolute value of lambda is negative [39,41]. Permutation tests of variable performance employed within the MaxEnt software platform used the training points to assess the relative contribution of each variable to the final averaged model in the context of the AUC statistic. A significant drop in the AUC statistic after a bioclimatic variable is removed suggests that the variable significantly contributes to the estimation of HS [43].

### Climate change and habitat suitability analysis

We projected HS for all 15 bumble bee species using bioclimatic data generated from three general circulation models (GCMs) with a 4.5 and 8.5 representative concentration pathway (RCP) for the year 2050 and 2070 [20]. The RCP is a greenhouse gas concentration trajectory that takes into account pollution and land-use change that occurred over the twenty-first century [20]. We elected to use an intermediate greenhouse emission scenario (RCP 4.5) and a high emission scenario (RCP 8.5) when projecting HS for each species in 2050 and 2070 in the Pacific Northwest. The three GCMs used in our analysis are the Community Climate System Model 4 (CCSM4), the Hadley Global Environmental Model 2-Atmosphere (HADGEM2-AO), and the Model for Interdisciplinary Research on Climate Earth System Model (MIROC-ESM-CHEM). The three GCMs were downloaded from the WorldClim database as described above (http://worldclim), and can be examined on the Climate Model Intercomparison Project Phase 5 (CMIP 5) (https://cmip.llnl.gov/).

SDMs for each species were averaged across the three GCMs according to RCP and year combinations to estimate HS under different climate change scenarios. To calculate HS change for each species, we subtracted projected HS based on the averaged GCM projections across the three models from contemporary HS estimates. We used a simple paired Wilcoxon test to determine if there was a significant difference in HS between contemporary and projected HS in 2050 and 2070. Except for the MaxEnt analysis, all statistical analyses were conducted with R v3.5.2 [38].

## Results

### Field Survey

In total, fifteen bumble bee species were detected in our survey. We captured 773 bumble bees across 23 unique field sites from 15 – 25 of July and 2 August 2013 (Table S1). Of the 773, 272 voucher specimens were curated and are currently housed at the NPIC in Logan, Utah (Table S2). The remaining 501 specimens not retained as vouchers were released at the collection site after field identification and tissue sampling. Average temperatures during the field survey were 22.3 ± 0.69 °C, average relative humidity was 50.6 ± 2.21% and average wind speed was 1.9 ± 0.41 kph. The total specimens surveyed from each park are EBLA = 35, FOVA = 55, LEWI = 33, MORA = 238, NOCA = 118, OLYM = 173, SAJH = 121 (Fig 1). The most abundant to least abundant species are as follows: *B. flavifrons* (*n* = 149), *B. sylvicola* (*n* = 119), *B. sitkensis* (*n* = 98), *B. bifarius* (*n* = 84), *B. mixtus* (*n* = 82), *B. melanopygus* (*n* = 69), *B. vosnesenskii* (*n* = 54), *B. rufocinctus* (*n* = 38), *B. caliginosus* (*n* = 24), *B. appositus* (*n* = 18), *B. californicus* (*n* = 14), *B. occidentalis* (*n* = 6), *B. vandykei* (*n* = 4), *B. griseocollis* (*n* = 1), *B. nevadensis* (*n* = 1), unidentified *Bombus* (*n* = 12). The unidentified *Bombus* included specimens that could not be reliably identified to species due to the poor condition of the physical characteristics needed for diagnosis [36]. Distribution and abundance of each species in the current study are available as supplementary figures (Figs S1-S15). *Bombus occidentalis* was detected at two sites in OLYM in the Royal Basin Area. This is the first time since 1955 that *B. occidentalis* has been detected within the boundaries of OLYM. However, it should be noted that a single *B. occidentalis* has been detected on Mt. Townsend in Olympic National Forest by a citizen scientist in 2011, and more recently in Seattle, Washington in 2013 [44]. All specimens identified in this survey are recorded in Table S1.

### Community analysis

To assess community richness and diversity, only specimens that were identified to species were used for the final analyses (*n* = 761). Thus, we removed the 12 unidentified specimens from the total 773 specimens surveyed. Across the sites assessed in our study, we found species richness to be positively correlated with rarefied species richness (*t* = 5.61, *df* = 17, *p* < 0.001, *r* = 0.81) and the inverse Simpson’s diversity index (*t* = 3.78, *df* = 17, *p* = 0.001, *r* = 0.68). Altitude was a significant predictor of species richness and diversity (simple linear regressions; richness: *F*_*1, 17*_ = 9.68, *p* = 0.01, *r*^*2*^ = 0.33; diversity: *F*_*1, 17*_ = 7.38, *p* = 0.01, *r*^*2*^ = 0.26) (Fig 2). Both species richness and diversity increased by 0.001 for each one meter increase in altitude [richness~3.24+0.001 (altitude), diversity~2.43+0.001 (altitude)]. Finally, NMDS analysis found altitude to be a significant predictor of community composition, with high and low altitude communities clearly demarcated (NMDS, *k* = 2, stress = 0.13, *r*^*2*^ = 0.66, *p* = 0.001). Specifically, bumble bee communities found at altitudes greater than 500 m shared species that were relatively unique to communities found at altitudes less than 500 m (Fig 3).

**Fig 2.**
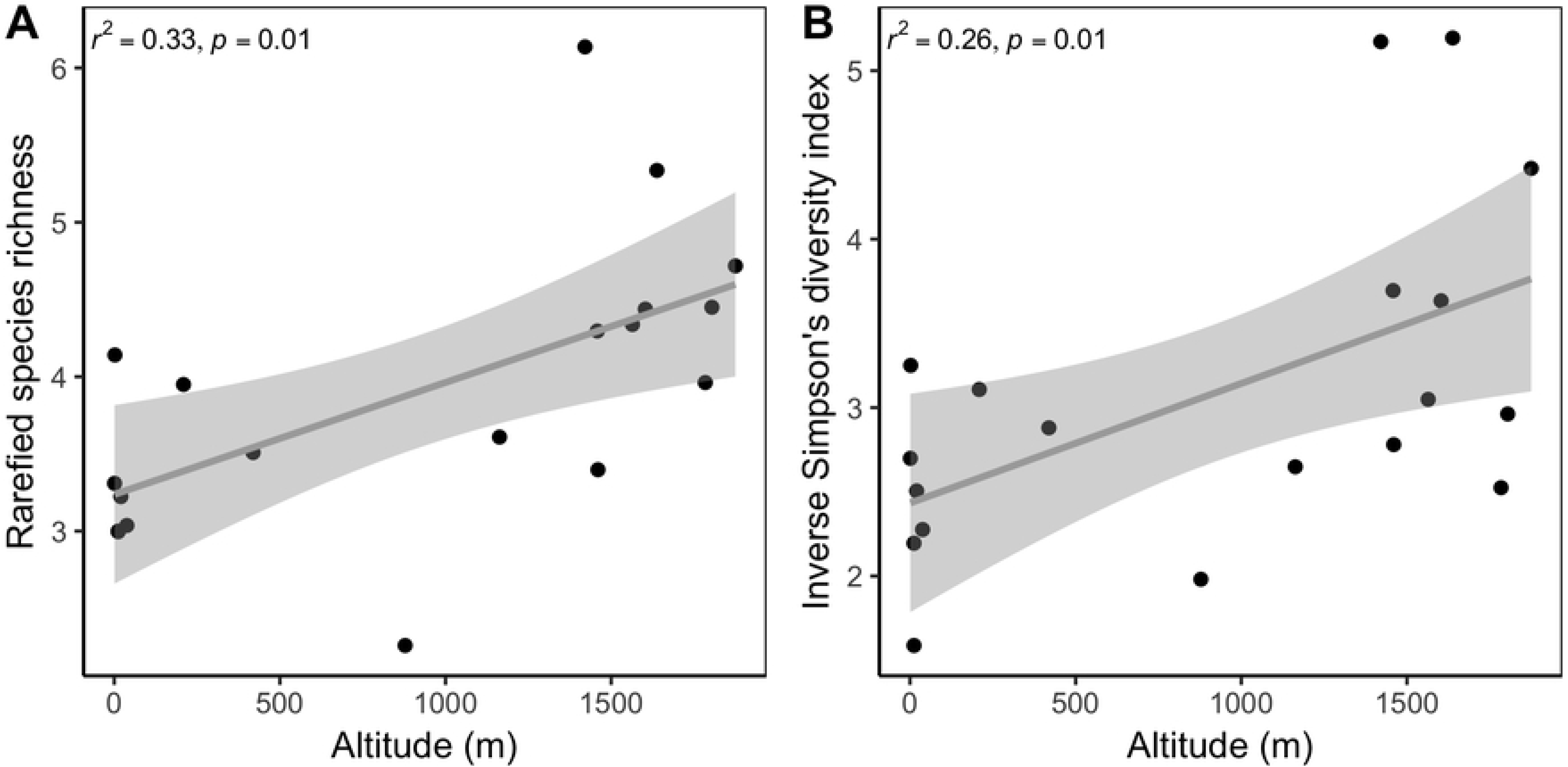
Distribution of rarified bumble bee species richness (A) and inverse Simpson’s diversity index (B) across an altitude gradient in US National Parks in the Pacific Northwest.

**Fig 3.**
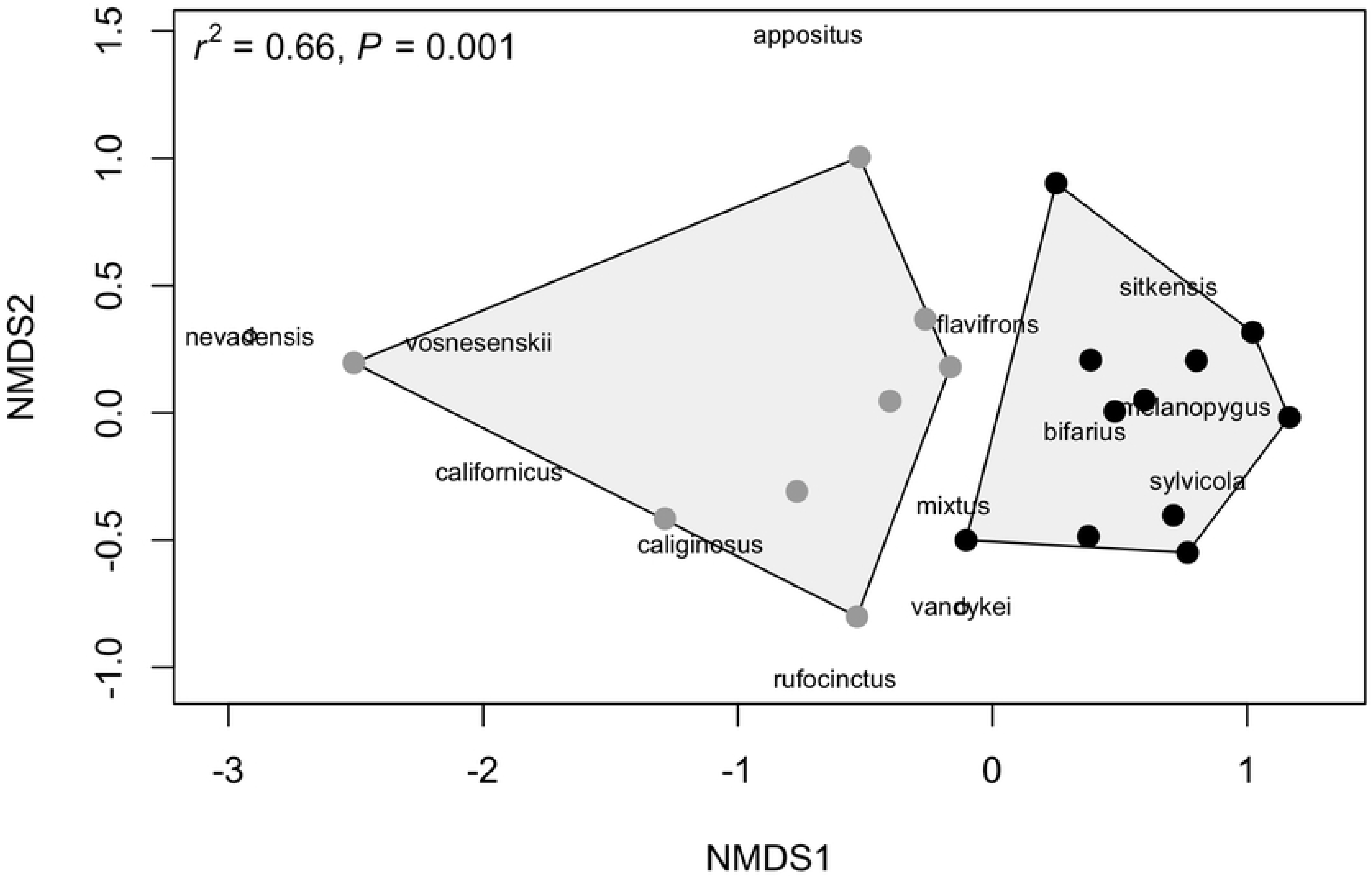
Nonmetric dimensional scaling analysis (NMDS) of bumble bee community dissimilarity across US National Parks in the Pacific Northwest. Locations clustered closer together suggest that bumble bee communities are more similar in composition. High altitude communities (gray points) are more similar in composition than low altitude communities (black points). Species names are presented in the figure to infer that species clustered closer together are found to co-occur, whereas species distributed further apart are less likely to co-occur. The point under *B. nevadensis* represents *B. griseocollis*, whereas the point under *B. vandykei* represents *B. occidentalis*.

### Species distribution modelling

We compiled a total of 113,551 specimens records across the 15 species assessed in this study from GBIF [45]. After filtering for unique spatial records, the dataset was reduced to 8,805 records. The average number of records per species available for each SDM is 587±129 SE. A summary of the number of records of the target species used for SDMs in our study is found in Table S3. Following correlation analysis, 11 of the 19 bioclimatic variables were used in the SDM: BIO1 = annual mean temperature, BIO2 = mean diurnal range (mean of monthly (maximum temp - minimum temp)), BIO3 = isothermality (BIO2/BIO7) (*100), BIO4 = temperature seasonality (standard deviation *100), BIO5 = maximum temperature of warmest month, BIO8 = mean temperature of wettest quarter, BIO9 = mean temperature of driest quarter, BIO13 = precipitation of wettest month, BIO14 = precipitation of driest month, BIO15 = precipitation seasonality (coefficient of variation), BIO18 = precipitation of warmest quarter. All 15 SDMs constructed in this study performed well, with AUC_test_ values between 0.79 and 0.96 (mean AUC_Test_: 0.87 ± 0.03) (Table S4).

Precipitation is a significant predictor of bumble bee HS across all 15 bumble bee species studied in the Pacific Northwest. Averaging all 15 species-specific SDM found that precipitation of wettest month (BIO13) contributed the most to SDM construction [18 ± 3.69 mean percent contribution on average plus/minus standard error (SE)], followed by mean temperature of wettest quarter (BIO 8) (17.4 ± 3.52 mean percent contribution on average), and precipitation of driest month (BIO14), (16.83 ± 5.02 mean percent contribution on average) (Table S5). Furthermore, when the bioclimatic variables are permuted in a SDM, BIO13 and BIO8 remain as important variables across the 15 different SDMs (BIO13: 20.46 ± 2. mean permutation importance; BIO 8: 14.31 ± 1.87 mean permutation importance), whereas precipitation seasonality (BIO15), was identified be the 2^nd^ most imported variable after permutation (16.02 ± 2.77 mean permutation importance).

### Climate change and habitat suitability analysis

Across both RCP scenarios and 2050 and 2070 time step combinations, it was clear that the vast majority of Pacific Northwest bumble bee species will undergo HS loss in the US National Parks within the study region (Fig 4) (Table 2). *Bombus vosnesenskii, B. sitkensis, B. caliginous*, and *B. californicus* might experience a small degree of HS gain within the study region (Fig 4). Relative to our sampled field sites in our study, *B. bifarius, B. flavifrons, B. melanopygus, B. mixtus*, and *B. sylvicola* are hypothesized to undergo significant HS loss in US National Park in the Pacific Northwest, whereas *B. vosnesenskii* and *B. sitkensis* are hypothesized to undergo significant HS gain by 2050 and 2070 (Paired Wilcoxon tests, all *P* < 0.05) (Fig 4) (Table S6). Finally, if species are to be prioritized by HS loss averaged across both RCP scenarios and time steps, the list of species from most vulnerable to least vulnerable to climate change are as follows: 1) *B. vandykei*, 2) *B. sylvicola*, 3) *B. bifarius*, 4) *B. melanopygus*, 5) *B. occidentalis*, 6) *B. flavifrons*, 7) *B. griseocollis*, 8) *B. nevadensis*, 9) *B. rufocinctus*, 10) *B. mixtus*, 11) *B. appositus*, 12) *B. sitkensis*, 13) *B. californicus*, 14) *B. caliginosus*, 15) *B. vosnesenskii* (Fig 5).

**Table 2.** Paired Wilcoxon (W) tests results comparing contemporary habitat suitability values and for the 4.5 and 8.5 representative concentration pathway (RCP) and future year scenarios (2050 and 2070) for 15 bumble bee species in US National Parks in the Pacific Northwest [20].

|  | Significant<br>at<br>P < 0.05? | RCP 4.5, Year 2050 |  | RCP 4.5, Year 2070 |  | RCP 8.5, Year 2050 |  | RCP 8.5, Year 2070 |  |
| --- | --- | --- | --- | --- | --- | --- | --- | --- | --- |
|  |  | W | P | W | P | W | P | W | P |
| *B. appositus* | NS | 4 | 0.333 | 4 | 0.333 | 4 | 0.333 | 4 | 0.333 |
| *B. bifarius* | * | 100 | 0.0002 | 100 | 0.0002 | 100 | 0.0002 | 100 | 0.0002 |
| *B. caliginosus* | NS | 9 | 0.1 | 9 | 0.1 | 9 | 0.1 | 9 | 0.1 |
| *B. californicus* | NS | 9 | 0.1 | 9 | 0.1 | 9 | 0.1 | 9 | 0.1 |
| *B. flavifrons* | * | 400 | 0.00001 | 400 | 0.00001 | 400 | 0.00001 | 400 | 0.00001 |
| *B. griseocollis* | NS | 1 | 1 | 1 | 1 | 1 | 1 | 1 | 1 |
| *B. melanopygus* | * | 169 | 0.0001 | 169 | 0.0001 | 169 | 0.0001 | 169 | 0.0001 |
| *B. mixtus* | * | 256 | 0.0001 | 256 | 0.0001 | 256 | 0.0001 | 256 | 0.0001 |
| *B. nevadensis* | NS | 1 | 1 | 1 | 1 | 1 | 1 | 1 | 1 |
| *B. occidentalis* | NS | 4 | 0.333 | 4 | 0.333 | 4 | 0.333 | 4 | 0.333 |
| *B. rufocinctus* | NS | 4 | 0.333 | 4 | 0.333 | 4 | 0.333 | 4 | 0.333 |
| *B. sitkensis* | * | 169 | 0.0001 | 169 | 0.0001 | 169 | 0.0001 | 169 | 0.0001 |
| *B. sylvicola* | * | 120 | 0.0001 | 121 | 0.00001 | 121 | 0.00001 | 120 | 0.0001 |
| *B. vandykei* | NS | 4 | 0.333 | 4 | 0.333 | 4 | 0.333 | 4 | 0.333 |
| *B. vosnesenskii* | * | 48 | 0.001 | 47 | 0.002 | 48 | 0.002 | 40 | 0.053 |

**Fig 4.**
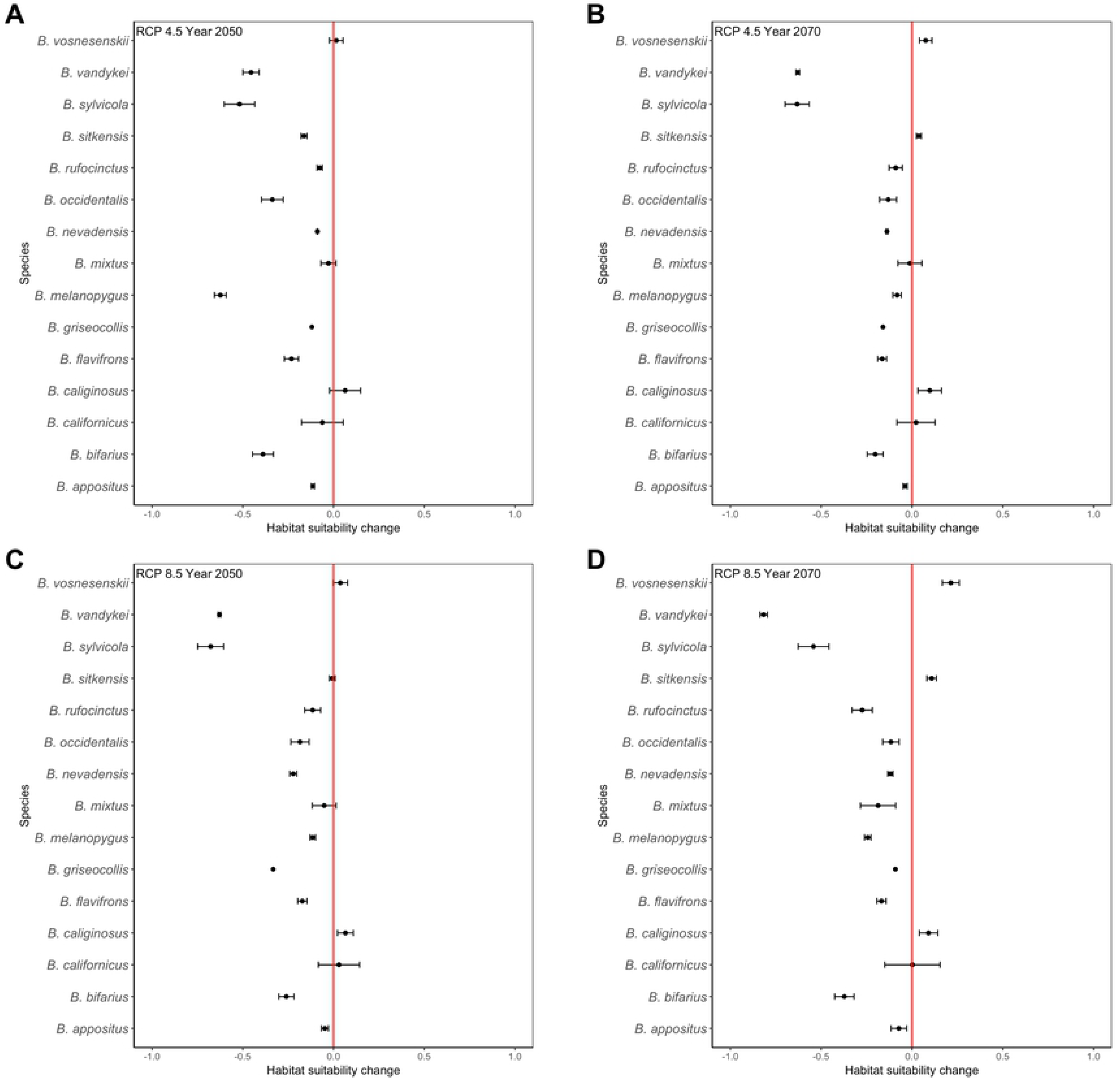
Habitat Suitability (HS) comparisons across 15 bumble bee species surveyed across and adjacent to US National Parks in the Pacific Northwest. Comparisons for each species are made between modeled HS of contemporary and future (2050 and 2070) distributions under two representative concentration pathways (RCP) [20]. (A) RCP 4.5, 2050, (B) RCP 4.5, 2070, (C) RCP 8.5, 2050, and (D) RCP 8.5, 2070. The X-axis represents the difference between contemporary and future HS. Values to the left of the dashed red line indicate a decrease in HS, whereas values to right of the dashed red line indicate an increase in HS.

**Fig 5.**
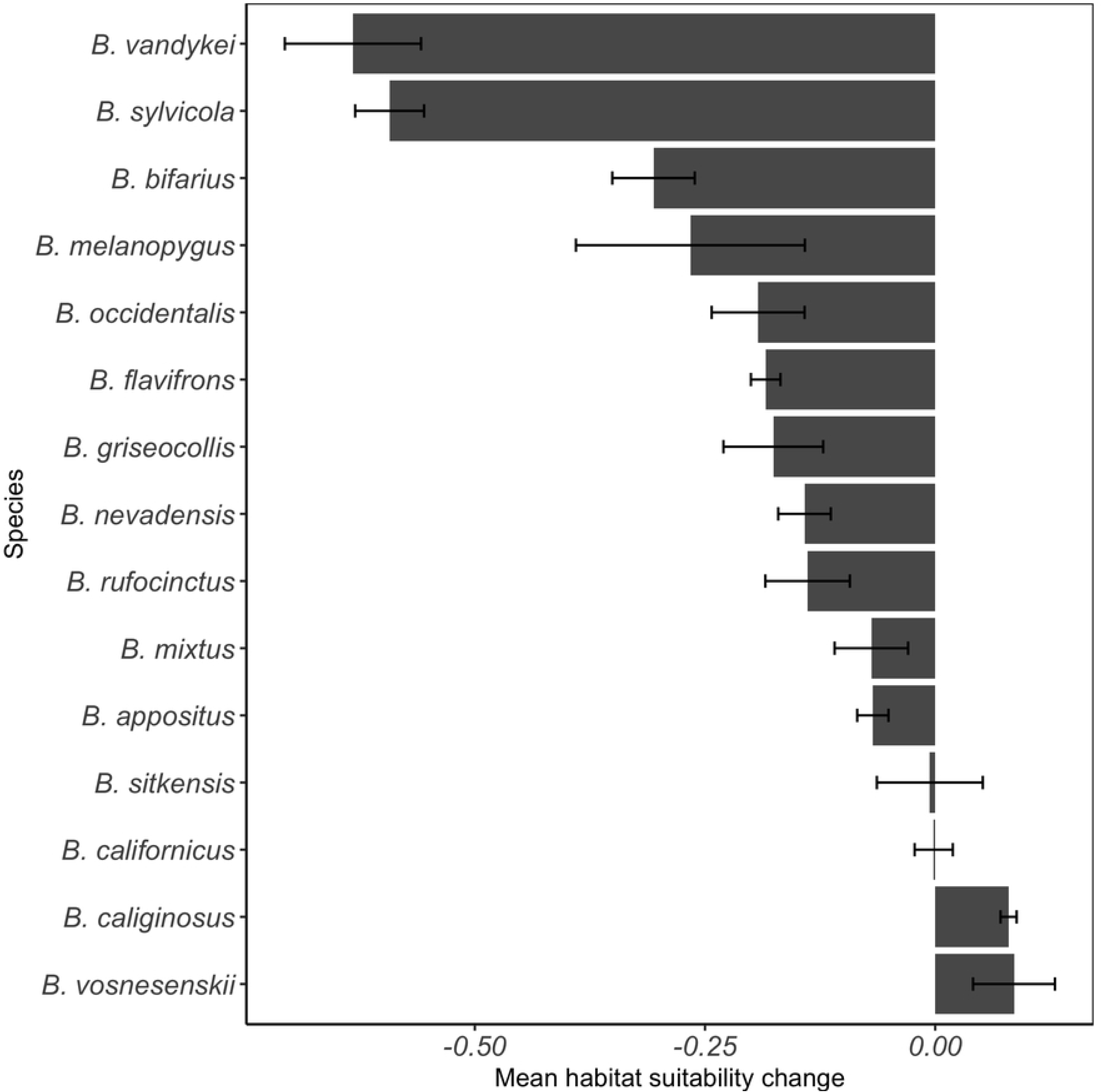
Mean (±SE) habitat suitability change across two relative concentration pathway scenarios (RCP 4.5 and 8.5) and two time steps (2050 and 2070) for 15 bumble bee species in US National Parks in the Pacific Northwest.

## Discussion

We discovered that bumble bee community composition and diversity can be predicted by their distribution across an altitude gradient in the Pacific Northwest. As expected, we found that both species richness and diversity were positively correlated with altitude (Fig 2). We also found that bumble bee community composition can be predicted by species’ distribution across an altitude gradient, with high altitude communities clustering differently than low altitude communities (Fig 3). Finally, an assessment of HS under two RCP scenarios (4.5 and 8.5) and two time steps (2050 and 2070) found that 80% of bumble bees found within the national park boundaries in the Pacific Northwest are projected to undergo HS loss (Figs 4, 5).

Our study supports the consensus that bumble bee community diversity and composition are predicted by bees’ distributions across an altitude gradient [15,46–49]. The greatest diversity of bumble bees in North America is found primarily in areas that are topographically complex environments, especially in mountainous regions of the western US [26,32]. Bumble bee species that are found predominantly in high alpine environments run the greatest risk of losing suitable habitat in the next 50 years. Why alpine bumble bees are most vulnerable to decline is likely due to the narrow bioclimatic niche they inhabit [22,50]. In our study, we find that bumble bee HS is best predicted by bioclimatic variables that capture precipitation estimates. The Pacific Northwest is a region of North America defined by rain forest as it receives a wealth of precipitation. The region is subject to receive more precipitation based on GCM projections for the region over the next 50 years [34], thus it is likely that that bumble bee HS will be impacted by changes in precipitation patterns in the region as precipitation is a significant predictor of bumble bee HS in our study.

The probable species composition of a bumble bee community can be demarcated based on altitude in the Pacific Northwest (Fig 3). In low altitude environments, the following bumble bees are likely to be detected: *B. nevadensis, B. griseocollis, B. vosnesenskii, B. californicus, B. caliginosus, B. appositus, B. rufocinctus*, and *B. flavifrons*. Alternatively the following bumble bees are likely to be detected in high altitude environments: *B. vandykei, B. occidentalis, B. mixtus, B. bifarius, B. sitkensis, B. melanopygus, B. sylvicola*. In our study, we found that 80% of the species studied are projected to experience significant HS loss regardless of the GCM RCP scenario or time step (Figs 4, 5). It is clear that high altitude bumble bee species will experience the greatest HS loss compared to low altitude bumble bee species (Fig 4).

Recent climate warming is suspected to have shifted bumble bee distributions across an altitude gradient, with low altitude environments losing species richness, and high altitude environments gaining species richness [15]. The shift in species richness is hypothesized to be an artifact of bumble bees dispersing to high altitudes as low altitude environments have become unsuitable bumble bee habitat. In the Pacific Northwest, bumble bee communities are more species rich and diverse in high altitude environments relative to low altitude environments (Fig 2). Therefore, if bumble bees from low altitude environments disperse to high altitude environments, it is likely that species will compete for floral, nest, and hibernacula resources in an environment that is also spatially limited in comparison to low altitude environments [51–53]. However, even if floral resources become limited, recent research suggests that some bumble bees might arguably be resilient to resource loss, as demonstrated *B. sylvicola* and *B. balteatus* populations in alpine environments of Colorado [16]. Selection for *B. sylvicola* and *B. balteatus* individuals with shorter proboscis to more effectively forage for floral resources has been documented in Colorado populations due to the decline of flowers with long corollas. In the case of Pacific Northwest bumble bees, the increase of competition by low altitude species coupled with expected shifts in floral resources abundance, diversity, and phenology might greatly impact the evolutionary trajectory of high altitude bumble bee species [16].

We discovered that *B. vandykei* will be the most vulnerable to climate change in the Pacific Northwest, as our models predicted that it will incur the greatest HS loss (63 ± 7 percent mean HS loss) (Fig 5). Historically, *B. vandykei* has not been detected on the Olympic Peninsula [36], and has only been recently detected within the Olympic Mountains of OLYM [29]. Furthermore, *B. vandykei* is a very rare bumble bee, and comprised only 0.52% (*n* = 4) of the total bumble bees collected in our survey (Fig S14). Bumble bees are well known to be misidentified due to convergent setal coloration patterns [36,54], thus, it is possible that the species may have been misidentified in previous assessments of the Pacific Northwest. Given that *B. vandykei* is a rare and potentially misidentified bumble bee, as evidence by lack of detection in historic surveys of the species [36], the classification of the species as most vulnerable to the effects of climate change is warranted.

Our HS analysis further suggests that *B. sylvicola* will experience great HS loss in the Pacific Northwest, with HS loss estimates between 52% and 67% under the different RCP scenario and year combinations (59 ± 4 percent mean HS loss) (Fig 5). Like *B. vandykei, B. sylvicola* has only been recently detected in OLYM [29], and yet it is poised to be one of the species most vulnerable to the effects of climate change. Populations of *B. sylvicola* in the Pacific Northwest form unique genetic clusters that are associated with their mountain province of origin, and are associated with low population genetic diversity [27]. Projected HS loss in the next 50 years coupled with low genetic diversity and isolation are factors that suggest that *B. sylvicola* is at great risk for population decline and extinction.

Finally, our survey found *B. occidentalis* to be restricted to high altitude environments based on the current sampling effort (Fig. 3). However, previous studies suggests that *B. occidentalis* was a historically abundant bumble bee species found at low altitude environments in the Pacific Northwest [10,55,56]. The hypothesized cause of decline in wild *B. occidentalis* is attributed to pathogens [10] and land-use change [7]. In our study, we did not assess pathogen vulnerability for all 15 bumble bee species. However, previous range-wide investigations of pathogen incidence in wild bumble bees suggest that several species that we documented in our study are associated with pathogens of concern including *Nosema bombi* and *Crithidia spp. [57]*. Future research could examine the intersection between climate and pathogen incidence in assessing bumble bee vulnerability in the Pacific Northwest [56].

Our study contributes to an important framework for identifying which bumble bee species in US National Parks are most vulnerable to projected climate change in the next 50 years [12]. Specifically, we categorize which bumble bee species are predicted to incur the greatest HS change in the Pacific Northwest. Bumble bees are poised to experience shifts in HS across both altitude and latitude in the next 50 years [3]. Species loss at low latitude environments and species gain in high latitude environments are estimated to occur predominantly eastern North America [2]. However, in western North America, where the landscape is characterized by a diversity of mountain ranges, the loss of bumble bee diversity is complex, likely due to differences in community assemblages across the region [2]. Along the Rocky Mountain spine, it appears that species gain is estimated to occur in some regions, likely due to a shift in HS across a latitude gradient. However, in all mountain provinces significant species loss in western North America is expected to occur across an altitude gradient [2]. Our regional study in the Pacific Northwest support the inference of Sirois-Delise and Kerr [2] that most bumble bee species will experience significant HS loss at low altitudes and latitudes, which will only be exacerbated by their inability to disperse to across geographic distance due to the lack of suitable habitat [27,28,43].

The results presented here will be useful in helping managers and stakeholders prioritize restoration and conservation efforts of bumble bees within US National Parks and adjacent areas in the Pacific Northwest. Specifically, as we have identified which species are most vulnerable to climate change, stakeholders can begin examining what types of other limiting factors might be useful to buffer the impacts of a warming climate on the most vulnerable. For example, stakeholders can provide adequate floral resources to the most vulnerable species by either protecting or planting species of critical importance [58]. Alternatively, combining SDM with population genetic data may inform the potential for habitat corridors as a mitigation strategy to ensure that vulnerable bumble bee species do not become isolated from adjacent populations [27,28]. Whatever the strategy, identifying which species is most vulnerable to climate change is a significant first step in the prioritization of conservation and management action.

## Acknowledgements

We thank Joyce Knoblett and Chris Lorion for their assistance with field surveys, Harold Ikerd for database support, and Thor Hanson for field survey assistance and hospitality. We also thank Dr. Jerry Freilich and Dr. Regina Rochefort for their unwavering support in the study of bumble bees in National Parks. This study was made possible with grant #NPS P13PG00149/FSN from the North Cascades and Coast Science Learning Network (NCCSLN) to JPS and WSS. JBK is currently supported by a David H. Smith Postdoctoral Fellowship. Permits to survey bumble bees from each US National Park are as follows: NOCA-2013-SCI-0012, SAJH-2013-SCI-0002, OLYM-2013-SCI-0049, OLYM-2014-SCI-0065, MORA-2013-SCI-0037, EBLA-2013-SCI-0001, LEWI-2013-SCI-0003, NCCO-2014-SCI-0006. A copy of the final report submitted to the US National Park Service may be found on the US Department of Interior, National Park Service Integrated Resource Management Application Portal, Research Permit and Reporting System at the following link: https://irma.nps.gov/rprs/IAR/Profile/103061 (Report #: NOCA-00139). The US Department of Agriculture, Agricultural Research Service (USDA-ARS) is an equal opportunity/affirmative action employer and all agency services are available without discrimination. Mention of commercial products and organizations in this manuscript is solely to provide specific information. It does not constitute endorsement by USDA-ARS over other products and organizations not mentioned.

## Supporting Information

**Table S1.** Database of the bumble bee specimens identified in the national parks of the North Coast and Cascades Network. Genus = genus, Species = species, M = male, F = female (Non-queen), Q = queen, Park = park acronym, Location Description = location description, Day0 = day, Mon0 = month, Year0 = year, Time0 = time survey started, Time1 = time survey ended, Floral host = flowering plant collected specimen on (if available), Col1 = collector 1, Col2 = collector 2, Col3 = collector 3, Col4 = collector 4, Temperature = temperature (degrees C), Wind Speed = wind speed in kph, Cloud cover = 1 (full cloud cover)/ 0 (full sun), Relative humidity = relative humidity.

**Table S2.** Voucher of the specimens collected at the national parks of the North Coast and Cascade Network following US National Park Service data deposition formatting. Catalog # = catalog number, Accession # = accession number, Cataloger = person who cataloged voucher, Class 1 = all Biology, Class 2 = all Animalia, Class 3 = all Insecta, Class 4 = all Hymenoptera, Collection Date = collection date, Collection # = not applicable, Collector = persons who collected specimens, County = county specimens collected, Elevation = elevation (m), Family = all Apoidea/Apidae, Identified by = species identification expert, Locality = location surveyed, Location = location where specimens are deposited, Obj/Science = species name, State = state code of locality, Habitat/Comm = not applicable, TRS = township and range search, Aspect = not applicable, Description = sex of specimen, if applicable.

**Table S3.** Distribution record summary of 15 bumble bees in USA (minimum longitude = −125; maximum longitude = −100; maximum latitude = 30; minimum latitude = 49; WGS 1984) queried from the Global Biodiversity Information Facility (GBIF; http://gbif.org). Records were used to construct species distribution models in the PNW (minimum longitude = −125; maximum longitude = −120; maximum latitude = 45; minimum latitude = 49; WGS 1984). Unique records = spatially unique records (duplicates removed from total GBIF records per species); Spatial filter (~5 km^2^) = spatially unique records are filtered to a resolution of ~5 km^2^; Proportion unique records = Unique records/Total GBIF records; Proportion unique & spatial filter (~5 km^2^) = Spatial filter (~5 km^2^)/Unique records.

**Table S4.** Area under the curve (AUC) species distribution model (SDM) performance summaries for 15 bumble bees species.

**Table S5.** Mean percent (%) contribution and permutation importance of 11 bioclimatic variables across 15 bumble bees species in US National Parks in the Pacific Northwest. Maximum = maximum mean value, Minimum = minimum mean value, SE = standard error. BIO1 = Annual Mean Temperature, BIO2 = Mean Diurnal Range (Mean of monthly (max temp - min temp)), BIO3 = Isothermality (BIO2/BIO7) (* 100), BIO4 = Temperature Seasonality (standard deviation *100), BIO5 = Max Temperature of Warmest Month, BIO8 = Mean Temperature of Wettest Quarter, BIO9 = Mean Temperature of Driest Quarter, BIO13 = Precipitation of Wettest Month, BIO14 = Precipitation of Driest Month, BIO15 = Precipitation Seasonality (Coefficient of Variation), BIO18 = Precipitation of Warmest Quarter.

**Figure S1.** Relative abundance of *B. appositus* across US National Parks in the Pacific Northwest.

**Figure S2.** Relative abundance of *B. bifarius* across US National Parks in the Pacific Northwest.

**Figure S3.** Relative abundance of *B. californicus* across US National Parks in the Pacific Northwest.

**Figure S4.** Relative abundance of *B. caliginosus* across US National Parks in the Pacific Northwest.

**Figure S5.** Relative abundance of *B. flavifrons* across US National Parks in the Pacific Northwest.

**Figure S6.** Relative abundance of *B. griseocollis* across US National Parks in the Pacific Northwest.

**Figure S7.** Relative abundance of *B. melanopygus* across US National Parks in the Pacific Northwest.

**Figure S8.** Relative abundance of *B. mixtus* across US National Parks in the Pacific Northwest.

**Figure S9.** Relative abundance of *B. nevadensis* across US National Parks in the Pacific Northwest.

**Figure S10.** Relative abundance of *B. occidentalis* across US National Parks in the Pacific Northwest.

**Figure S11.** Relative abundance of *B. rufocinctus* across US National Parks in the Pacific Northwest.

**Figure S12.** Relative abundance of *B. sitkensis* across US National Parks in the Pacific Northwest.

**Figure S13.** Relative abundance of *B. sylvicola* across US National Parks in the Pacific Northwest.

**Figure S14.** Relative abundance of *B. vandykei* across US National Parks in the Pacific Northwest.

**Figure S15.** Relative abundance of *B. vosnesenskii* across US National Parks in the Pacific Northwest.

## References

1. Winfree R, Aguilar R, Vázquez DP, LeBuhn G, Aizen MA. A meta-analysis of bees’ responses to anthropogenic disturbance. Ecology. 2009;90: 2068–2076.

2. Sirois-Delisle C, Kerr JT. Climate change-driven range losses among bumblebee species are poised to accelerate. Sci Rep. 2018;8: 14464.

3. Kerr JT, Pindar A, Galpern P, Packer L, Potts SG, Roberts SM, et al. Climate change impacts on bumblebees converge across continents. Science. 2015;349: 177–180.

4. Bartomeus I, Ascher JS, Wagner D, Danforth BN, Colla S, Kornbluth S, et al. Climate-associated phenological advances in bee pollinators and bee-pollinated plants. Proc Natl Acad Sci USA. 2011;108: 20645–20649.

5. Goulson D, Lye GC, Darvill B. Decline and conservation of bumble bees. Annu Rev Entomol. 2008;53: 191–208.

6. Cameron SA, Lim HC, Lozier JD, Duennes MA, Thorp R. Test of the invasive pathogen hypothesis of bumble bee decline in North America. Proc Natl Acad Sci USA. 2016; doi:10.1073/pnas.1525266113.

7. McFrederick QS, LeBuhn G. Are urban parks refuges for bumble bees *Bombus* spp. (Hymenoptera: Apidae)? Biol Conserv. 2006;129: 372–382.

8. Herrera JM, Ploquin EF, Rodríguez-Pérez J, Obeso JR. Determining habitat suitability for bumblebees in a mountain system: a baseline approach for testing the impact of climate change on the occurrence and abundance of species. J Biogeogr. 2014;41: 700–712.

9. Sachman-Ruiz B, Narváez-Padilla V, Reynaud E. Commercial *Bombus impatiens* as reservoirs of emerging infectious diseases in central México. Biol Invasions. 2015; 1–11.

10. Cameron SA, Lozier JD, Strange JP, Koch JB, Cordes N, Solter LF, et al. Patterns of widespread decline in North American bumble bees. Proc Natl Acad Sci USA. 2011;108: 662–667.

11. Whitehorn PR, O’Connor S, Wackers FL, Goulson D. Neonicotinoid pesticide reduces bumble bee colony growth and queen production. Science. 2012;336: 351–352.

12. Williams SE, Shoo LP, Isaac JL, Hoffmann AA, Langham G. Towards an integrated framework for assessing the vulnerability of species to climate change. PLoS Biol. 2008;6: 2621–2626.

13. Foden WB, Butchart SHM, Stuart SN, Vié J-C, Akçakaya HR, Angulo A, et al. Identifying the world’s most climate change vulnerable species: a systematic trait-based assessment of all birds, amphibians and corals. PLoS One. 2013;8: e65427.

14. Rasmann S, Pellissier L, Defossez E, Jactel H, Kunstler G. Climate-driven change in plant--insect interactions along elevation gradients. Funct Ecol. 2014;28: 46–54.

15. Fourcade Y, Åström S, Öckinger E. Climate and land-cover change alter bumblebee species richness and community composition in subalpine areas. Biodivers Conserv. 2019;28: 639– 653.

16. Miller-Struttmann NE, Geib JC, Franklin JD, Kevan PG, Holdo RM, Ebert-May D, et al. Functional mismatch in a bumble bee pollination mutualism under climate change. Science. 2015;349: 1541–1544.

17. Dirnböck T, Essl F, Rabitsch W. Disproportional risk for habitat loss of high-altitude endemic species under climate change. Glob Chang Biol. 2011;17: 990–996.

18. Parmesan C. Ecological and evolutionary responses to recent climate change. Annu Rev Ecol Evol Syst. 2006;37: 637–669.

19. Williams PH, Osborne JL. Bumblebee vulnerability and conservation world-wide. Apidologie. 2009;40: 367–387.

20. IPCC. Climate Change 2014: Synthesis Report. Contribution of Working Groups I, II and III to the Fifth Assessment Report of the Intergovernmental Panel on Climate Change. 2014; 151.

21. Hoffmann AA, Willi Y. Detecting genetic responses to environmental change. Nat Rev Genet. 2008;9: 421–432.

22. Williams P, Colla S, Xie Z. Bumblebee vulnerability: common correlates of winners and losers across three continents. Conserv Biol. 2009;23: 931–940.

23. Hoffmann AA, Sgrò CM. Climate change and evolutionary adaptation. Nature. 2011;470: 479–485.

24. Rubidge EM, Patton JL, Lim M, Cole Burton A, Brashares JS, Moritz C. Climate-induced range contraction drives genetic erosion in an alpine mammal. Nat Clim Chang. 2012;2: 285–288.

25. Williams P. Bumblebees of the World. In: The Trustees of The Natural History Museum, London [Internet]. 2017 [cited 8 May 2017]. Available: http://www.nhm.ac.uk/research-curation/research/projects/bombus/

26. Koch JB, Strange JP, Williams P. Bumble Bees of the Western United States. San Francisco, CA: Pollinator Partnership; 2012.

27. Koch JB, Looney C, Sheppard WS, Strange JP. Patterns of population genetic structure and diversity across bumble bee communities in the Pacific Northwest. Conserv Genet. Springer; 2017;18: 507–520.

28. Lozier JD, Strange JP, Koch JB. Landscape heterogeneity predicts gene flow in a widespread polymorphic bumble bee, *Bombus bifarius* (Hymenoptera: Apidae). Conserv Genet. Springer; 2013;14: 1099–1110.

29. Koch JB, Looney C, Sheppard WS, Strange JP. Range extension of two bumble bee species (Hymenoptera: Apidae) into Olympic National Park. Northwest Sci. 2016;90: 228–234.

30. Lozier JD, Jackson JM, Dillon ME, Strange JP. Population genomics of divergence among extreme and intermediate color forms in a polymorphic insect. Ecol Evol. 2016;6: 1075– 1091.

31. Fraser DF, Copley CR, Elle E. Changes in the status and distribution of the Yellow-faced Bumble Bee. J Entomol Soc British Columbia. 2012: 31–37.

32. Strange JP, Tripodi AD. Characterizing bumble bee (*Bombu*s) communities in the United States and assessing a conservation monitoring method. Ecol Evol. 2019;00: 1.

33. Koch JB, Lozier J, Strange JP, Ikerd H, Griswold T, Cordes N, et al. USBombus, a database of contemporary survey data for North American Bumble Bees (Hymenoptera, Apidae, *Bombus*) distributed in the United States. Biodivers Data J. 2015; e6833.

34. Mote PW, Salathé EP Jr. Future climate in the Pacific Northwest. Clim Change. 2010;102: 29–50.

35. Pacifici M, Foden WB, Visconti P, Watson JEM, Butchart SHM, Kovacs KM, et al. Assessing species vulnerability to climate change. Nat Clim Chang. 2015;5: 215.

36. Thorp R, Horning DS Jr, Dunning LL. Bumble Bees and Cuckoo Bumble Bees of California (Hymenoptera: Apidae). Berkeley and Los Angeles, CA: University of California Press; 1983.

37. Gotelli NJ. A primer of Ecology. Sunderland, MA: Sinauer Associates, Inc.; 2008.

38. R Core Development Team. R: A language and environment for statistical computing. R Foundation for Statistical Computing [Internet]. Vienna, Austria; 2018. Available: http://www.R-project.org

39. Phillips SJ, Dudík M, Schapire RE. A Maximum Entropy Approach to Species Distribution Modeling. Proceedings of the Twenty-first International Conference on Machine Learning. New York, NY, USA: ACM; 2004. p. 83.

40. Phillips SJ, Anderson RP, Dudík M, Schapire RE, Blair ME. Opening the black box: an open-source release of Maxent. Ecography. 2017;40: 887–893.

41. Elith J, Phillips SJ, Hastie T, Dudík M, Chee YE, Yates CJ. A statistical explanation of MaxEnt for ecologists. Divers Distrib. 2011;17: 43–57.

42. Hijmans RJ, Cameron SE, Parra JL, Jones PG, Jarvis A. Very high resolution interpolated climate surfaces for global land areas. Int J Climatol. 2005;25: 1965–1978.

43. Koch JB, Vandame R, Mérida-Rivas J, Sagot P, Strange J. Quaternary climate instability is correlated with patterns of population genetic variability in *Bombus huntii*. Ecol Evol. 2018;108: 20645.

44. Doughton S. Native bee species spotted for first time since ’90s. The Seattle Times. 14 Jul 2013. Available: https://www.seattletimes.com/seattle-news/native-bee-species-spotted-for-first-time-since-rsquo90s/

45. GBIF.org. GBIF Occurrence Download [Internet]. Available: https://doi.org/10.15468/dl.nwko8h

46. Hatfield RG, LeBuhn G. Patch and landscape factors shape community assemblage of bumble bees, *Bombus* spp. (Hymenoptera: Apidae), in montane meadows. Biol Conserv. 2007;139: 150–158.

47. Williams P, Tang Y, Yao J, Cameron S. The bumblebees of Sichuan (Hymenoptera: Apidae, Bombini). System Biodivers. 2009;7: 101–189.

48. Marshall L, Biesmeijer JC, Rasmont P, Vereecken NJ, Dvorak L, Fitzpatrick U, et al. The interplay of climate and land use change affects the distribution of EU bumblebees. Glob Chang Biol. 2018;24: 101–116.

49. Hofmann MM, Fleischmann A, Renner SS. Correction to: Changes in the bee fauna of a German botanical garden between 1997 and 2017, attributable to climate warming, not other parameters. Oecologia. 2018; 187: 701–706.

50. Williams PH, Araújo MB, Rasmont P. Can vulnerability among British bumblebee (Bombus) species be explained by niche position and breadth? Biol Conserv. 2007;138: 493–505.

51. Ings TC, Ward NL, Chittka L. Can commercially imported bumble bees out-compete their native conspecifics? J Appl Ecol. 2006;43: 940–948.

52. Ishii HS, Kadoya T, Kikuchi R, Suda S-I, Washitani I. Habitat and flower resource partitioning by an exotic and three native bumble bees in central Hokkaido, Japan. Biol Conserv. 2008;141: 2597–2607.

53. Russo L. Positive and Negative impacts of non-native bee species around the world. Insects. 2016;7. doi:10.3390/insects7040069.

54. Koch JB. The decline and conservation status of North American bumble bees. digitalcommons.usu.edu; 2011; Available: https://digitalcommons.usu.edu/etd/1015/.

55. Koch JB, Strange JP. Constructing a species database and historic range maps for North American Bumble bees (*Bombus* sensu stricto Latrielle) to inform conservation decisions. Uludag Bee Journal. 2009;9: 97–108.

56. Rao S, Stephen WP, Kimoto C, DeBano SJ. The status of the ‘Red-Listed’ *Bombus occidentalis* (Hymenoptera: Apiformes) in Northeastern Oregon. sNorthwest Sci. 2011;85: 64–67.

57. Cordes N, Huang W-F, Strange JP, Cameron SA, Griswold TL, Lozier JD, et al. Interspecific geographic distribution and variation of the pathogens Nosema bombi and Crithidia species in United States bumble bee populations. J Invertebr Pathol. 2012;109: 209–216.

58. Potts SG, Vulliamy B, Dafni A, Ne’eman G, Willmer P. Linking bees and flowers: How do floral communities structure pollinator communities? Ecology. 2003;84: 2628–2642.

